# Environments and host genetics influence the geographic distribution of plant microbiome structure

**DOI:** 10.1101/2023.03.20.533563

**Authors:** Na Wei, Jiaqi Tan

## Abstract

1. To understand how microbiota influence plant populations in nature, it is important to examine the geographic distribution of plant-associated microbiomes and the underlying mechanisms. However, we currently lack a fundamental understanding of the biogeography of plant microbiomes and the environmental and host genetic factors that shape their distribution.
2. Leveraging the broad distribution and extensive genetic variation in duckweeds (the *Lemna* species complex), we identified the key factors that influenced the geographic distribution of plant microbiome diversity and compositional variation.
3. In line with the pattern observed in microbial biogeography based on free-living environmental microbiomes, we observed higher bacterial richness in temperate regions relative to lower latitudes in duckweed microbiomes (with 10% higher in temperate populations). Our analyses revealed that temperature and sodium concentration in aquatic environments had a negative impact on duckweed bacterial richness, whereas temperature, precipitation, pH, and concentrations of phosphorus and calcium, along with duckweed genetic variation, influenced the geographic variation of duckweed bacterial community composition.
4. The findings add significantly to our understanding of host-associated microbial biogeography and provide insights into the relative impact of different ecological processes, such as selection by environments and host genetics, dispersal, and chance, on plant microbiome assembly. These insights have important implications for predicting plant microbiome vulnerability and resilience under changing climates and intensifying anthropogenic activities.

## Introduction

Plants host diverse microbial symbionts, and these microbial symbionts are important for the functioning of plants within ecosystems (Laforest-Lapointe et al., 2017; Tan et al., 2023). To better understand the influence of microbiomes on plant populations across geographic ranges in nature, it is important to examine the distribution patterns of plant-associated microbiomes and the mechanisms that drive these patterns. While our knowledge of microbial biogeography has advanced greatly through investigating free-living environmental microbiomes across terrestrial, marine, and atmospheric ecosystems (Tedersoo et al., 2014; Sunagawa et al., 2015; Bahram et al., 2018; Zhao et al., 2022), significant knowledge gaps exist as to what drives the geographic distribution of local microbiome diversity and compositional variation across populations in host-associated microbiomes, such as plant microbiomes. It is also unclear whether the principles of microbial biogeography derived from free-living microbiomes can be generalized to host-associated microbiomes.

While various biogeography theories have been proposed to explain the distribution of diversity in plants and animals (Rosenzweig, 1995), microbial diversity does not always follow the same patterns as observed in their macroscopic counterparts (Chu et al., 2020). For instance, fungal diversity in soil microbiomes follows a latitudinal gradient, decreasing from lower to higher latitudes (Tedersoo et al., 2014; Bahram et al., 2018), similar to the patterns observed in plants and animals (Rosenzweig, 1995). However, global bacterial diversity peaks in temperate regions across soil, marine, and airborne microbiomes (Tedersoo et al., 2014; Sunagawa et al., 2015; Bahram et al., 2018; Zhao et al., 2022). The biogeography of free-living environmental microbiomes, therefore, indicates that ecological factors that may or may not follow latitudinal gradients can drive the geographic distribution of microbial diversity. Factors that exhibit a correlation with latitude may contribute to an observed latitudinal gradient of microbial diversity, as seen in the case of precipitation which predicts the distribution of soil fungal richness (Tedersoo et al., 2014). By contrast, factors that do not exhibit such a correlation may weaken and lead to a distinct biogeographic pattern, as seen in the case of pH which predicts the distribution of soil bacterial richness (Fierer & Jackson, 2006; Bahram et al., 2018). Compared to free-living microbiomes, plant microbiomes are subject to host-imposed niche filtering (Wei & Ashman 2018; Wei et al., 2022), which has the potential to reinforce or modify the role of environmental factors in driving microbial biogeography. The extent to which host plants, such as their genetic variation, affect the geographic distribution of microbial diversity may depend on whether hosts have adapted to the same or different environmental factors that influence microbial diversity. If hosts exhibit adaptation to the same environmental factors as microbes, host genetic variation may contribute to the observed patterns of microbial diversity caused by environments, while dissimilar adaptations may weaken the patterns.

Another notable pattern of microbial biogeography is the decay in microbial community similarity over geographic distance. Such distance decay is common across ecosystems (Sunagawa et al., 2015; Bahram et al., 2018; Zhao et al., 2022), and can arise due to a combination of ecological processes including dispersal limitation, environmental heterogeneity, and chance (Vellend, 2010; Mittelbach & Schemske, 2015). While dispersal limitation and chance promote stochasticity and play a major role in driving the geographic variation of microbial community composition in nature (Zhao et al., 2022), environmental heterogeneity is also important and drives niche-based selection (Fierer & Jackson, 2006; Tedersoo et al., 2014; Sunagawa et al., 2015; Bahram et al., 2018). For instance, in terrestrial ecosystems, variation in soil pH and nutrient concentration leads to variation in soil bacterial community composition (Fierer & Jackson, 2006; Bahram et al., 2018). Similarly, in marine ecosystems, temperature variation is the primary driver of variation in bacterial community composition in surface waters (Sunagawa et al., 2015). In addition to selection by environments, selection by host genetic variation may also contribute to the geographic variation of microbiome composition associated with plants, and the respective and collective roles of host genetic and environmental variation will depend on the extent to which host genetic variation is shaped by the same or different environmental factors.

To enhance our understanding of the geographic distribution of microbiome diversity and compositional variation in plant microbiomes and the underlying mechanisms, we leveraged the broad distribution and extensive genetic variation of the duckweed, *Lemna* species complex (referred to as *Lemna* or duckweeds for simplicity). *Lemna* is floating aquatic plants commonly found in slow-moving freshwater ecosystems worldwide (Landolt, 1986), and plays an important role in ecosystem functions and services, such as carbon sequestration, phytoremediation, biofuel production, and animal feedstock (Cao et al., 2018; Acosta et al., 2021). In *Lemna*, hybridization has led to extensive genetic variation, making this species complex morphologically similar (Braglia et al., 2021). In this study, we examined *Lemna* microbiomes across 34 different populations in the United States, covering both the cool temperate and hot humid subtropical regions. Our purposes were twofold. First, we sought to test the hypothesis that bacterial richness is higher in temperate regions relative to lower latitudes and uncover the environmental and host genetic factors driving the observed pattern. Second, we aimed to quantify the respective impact of ecological processes (e.g. selection, dispersal limitation, chance) on microbiome assembly and identify the environmental and host genetic factors driving the geographic variation of bacterial community composition.

## Materials and Methods

### Field collection

We collected *Lemna* and its microbiomes from 34 populations in the northern and southern range of its distribution in the United States (Fig. 1a and Table S1): Ohio (OH, Cleveland, *N* = 8; Columbus, *N* = 5), New Hampshire (NH, *N* = 2), Massachusetts (MA, *N* = 2), Rhode Island (RI, *N* = 2), Louisiana (LA, *N* = 7), Georgia (GA, *N* = 4), and South Carolina (SC, *N* = 4). The field sampling was conducted during the fast-growing season of duckweeds during June–August 2022. In addition, we collected samples from the two Massachusetts populations during the late growing season in October 2022 to confirm the negligible influence of temporary dynamics on duckweed microbiomes, relative to the other factors we investigated in this study. Specifically, at each population, we collected duckweeds using ethanol-sterilized forks into sterile plastic bags and stored them at 4 ºC until microbiome isolation within five days. We also measured the pH, conductivity (EC), and total dissolved solids (TDS) of the aquatic environment at each population using an Ohaus ST20M-B meter (Ohaus Corporation, Parsippany, New Jersey). Additionally, we collected 100 mL surface water in sterile centrifuge tubes and sent to the Wetland Biochemistry Analytical Services at Louisiana State University for additional water chemistry analysis (total organic carbon, TOC; total nitrogen, TN; total phosphorus, TP; major and trace elements including Na, Ca, Mg, Fe, Si, Cu, Zn, Mn, Pb, Cd; Table S1).

**Fig. 1.**
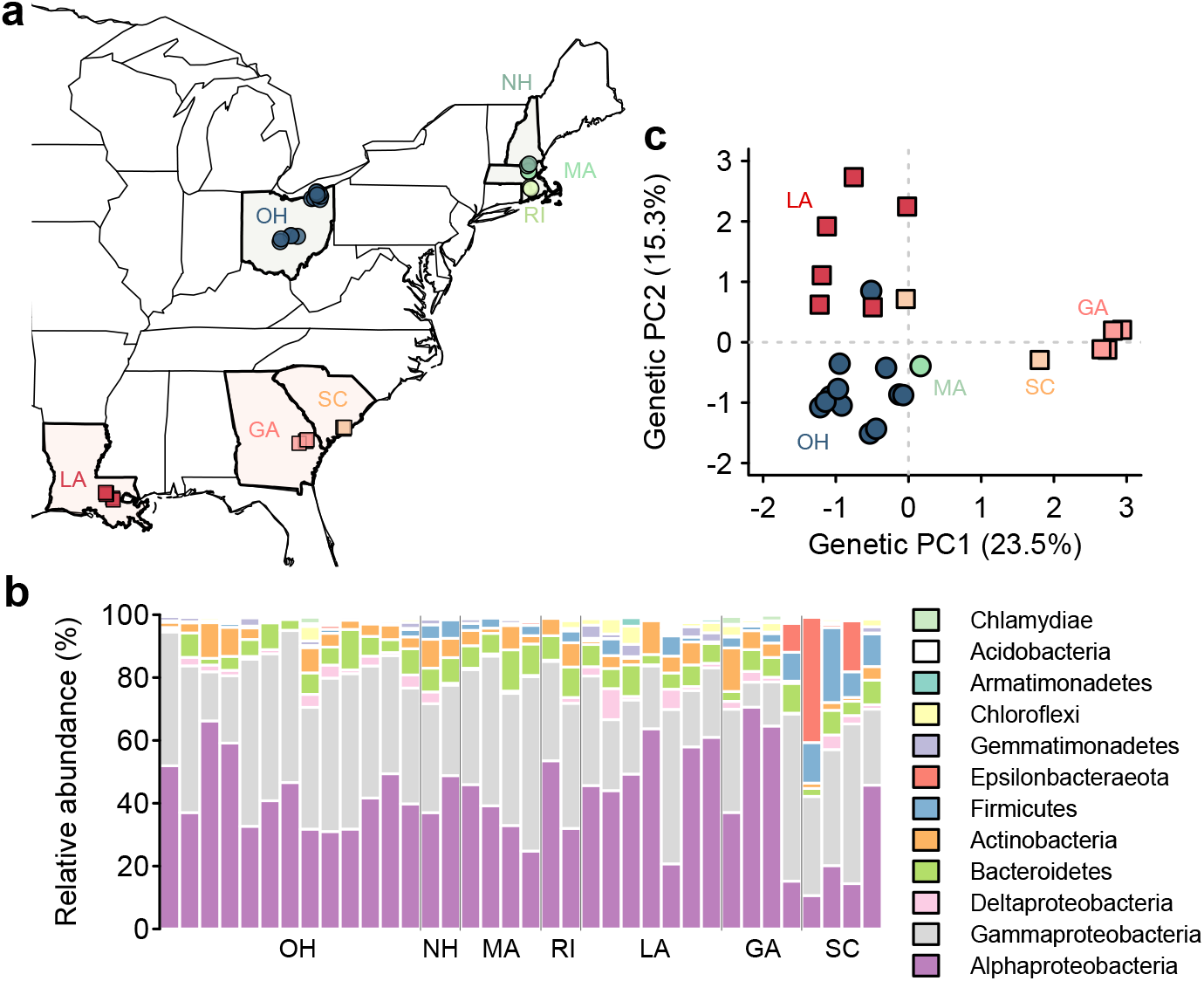
*Lemna* populations and microbiomes. (a) We collected the *Lemna* species complex from the northern and southern range of its distribution in the United States (34 total populations: OH, 13; NH, 2; MA, 2; RI, 2; LA, 7; GA, 4; SC, 4). (b) The top 10 most abundant phyla (class level for Proteobacteria) of *Lemna* bacterial microbiomes. The two MA populations (referred to as MA.1 and MA.2) were sampled at two separate times during the peak (June–August) and the end of the growing reason (October) in 2022. The order of the four MA samples in the plot follows MA.1 (peak and end season) and then MA.2 (peak and end season). (c) We obtained *Lemna* genetic data for 25 out of the 34 populations based on ISSR markers. *Lemna* genetic variation was examined using a PCA.

### Microbiome isolation and sequencing

Duckweed microbiome isolation was conducted sterilely under a laminar flow hood. For each population, we used sterilized forceps to remove debris from *Lemna*, and rinsed *c*. 500 individuals in 20 mL sterile water to remove environmental microbes from their aquatic habitats. These individual plants were then transferred to 20 mL sterile 0.25× phosphate buffered saline. We collected epiphytic microbiomes by vortexing for 20 min, sonicating at 40 kHz for 5 min, and centrifuging at 13,200 rpm for 10 min. Microbial cells (from 5 mL out of the 20 mL epiphytic microbiome wash) were used for DNA extraction using cetyltrimethylammonium bromide (CTAB) and purified using polyethylene glycol (PEG) 8000. Briefly, microbial pellets were lysed with 500 µL sterile CTAB buffer (2% w/v CTAB, 100 mM Tris-HCl, 20 mM EDTA, 1.4 M NaCl, 5 mM ascorbic acid, and 10 mM dithiothreitol) and two autoclaved 4 mm stainless steel beads on a Vortex Genie 2 (Scientific Industries, Bohemia, New York) for 40 min. An equal volume (500 µL) of chloroform : isoamyl alcohol (24 : 1) was then added for phage separation at 13,200 rpm for 5 min. DNA was then recovered by adding the upper phase to 1 mL of cold pure ethanol overnight at -20ºC and centrifuging at 13,200 rpm for 5 min. Pelleted DNA was washed with 500 µL of cold 70% ethanol and eluted in sterile TE buffer. We further purified the eluted DNA by conducting an additional round of chloroform : isoamyl alcohol phase separation, and then DNA was recovered by adding the upper phase to an equal volume of autoclaved PEG 8000 (20% w/v PEG 8000, 2.5 M NaCl), incubating at 37 ºC for 30 min, and centrifuging at 13,200 rpm for 5 min. Purified DNA pellet was washed with cold 70% ethanol and eluted in 60 µL sterile TE buffer and sent to the Argonne National Laboratory for bacterial library preparation (16S rRNA V5–V6 region, 799f–1115r primer pair) and sequencing using Illumina MiSeq (paired-end 250 bp).

The paired-end (PE) reads were used for detecting bacterial amplicon sequence variants (ASVs) using the package DADA2 v1.20.0 (Callahan et al., 2016) in R v4.1.0 (R Core Team, 2021). Following previous pipelines (Wei et al., 2021, 2022), the PE reads were trimmed and quality filtered [truncLen = c(240, 230), trimLeft=c(10, 0), maxN = 0, truncQ = 2, maxEE = c(2,2)] and then used for unique sequence identification that took into account sequence errors. The PE reads were then end joined (minOverlap = 20, maxMismatch = 4) for ASV detection and chimera removal. The ASVs were assigned with taxonomic identification based on the SILVA reference database (132 release NR 99) implemented in DADA2. The ASVs were further filtered before conversion into a bacterial community matrix using the package phyloseq (McMurdie & Holmes, 2013). First, we removed non-focal ASVs (Archaea, chloroplasts, and mitochondria). Second, we conducted rarefaction analysis using the package iNEXT (Hsieh et al., 2020) to confirm that the sequencing effort was sufficient to capture duckweed bacterial richness (Fig. S1). We further normalized per-sample reads (median = 20,192 reads) by rarefying to 10,000 reads. Three populations with fewer reads (one from OH: 9787 reads; two from GA: 5775 and 9484 reads, respectively) were normalized to 10,000 reads following the previous pipeline (Wei et al., 2021). Lastly, we removed low-frequency ASVs (<0.001% of total observations). The final bacterial community matrix consisted of 4880 ASVs across the 36 samples from 34 different populations.

### Lemna genotyping

After microbiome isolation, duckweeds were bleached to create axenic plants. Briefly, *c*. 30 clusters (100 plants) per population were bleached in 15 mL 1% sodium hypochlorite until clusters turned white, and then washed in 15 mL sterile water three times. Individual clusters were then grown in 0.5× Hoagland salt (PhytoTech Labs, Lenexa, Kansas) with 0.5% sucrose under 24 ºC and 16 h light for contamination check. A single axenic cluster was selected from a population (referred to as one genetic line) for further propagation in the same media for DNA extraction. Fresh duckweeds (*c*. 60 clonal plants) of each genetic line were used for DNA extraction using E.Z.N.A. SP Plant DNA Kit (Omega Bio-Tek Inc., Norcross, Georgia) and eluted in 100 µL sterile TE buffer. To examine duckweed genetic variation, we genotyped the genetic lines (*N* = 25, due to the unsuccess in generating some of the axenic genetic lines; Table S2). We used three polymorphic ISSR markers (UBC827, UBC855, UBC856) that generated a total of 46 polymorphic bands across the genetic lines (Table S2). PCRs were carried out in 10-µL reactions that contained 1.5 µL of extracted DNA, 0.5 µM primer, 4 mM MgCl_2_, 0.5 mg/mL BSA, 5 μL GoTaq Colorless Master Mix (Promega Corporation, Madison, Wisconsin) including 200 μM of each dNTP and 1 unit Taq DNA polymerase, and H_2_O. PCRs followed a standard protocol: 94 ºC for 5 min; 40 cycles of 94 ºC for 1 min, 52 ºC for 1 min, and 72 ºC for 1 min; and a final extension at 72 ºC for 5 min. PCR amplicons were quantified with GeneRuler 100bp plus DNA Ladder (Thermo Fisher Scientific Inc., Waltham, Massachusetts) on 1.5% agarose gels in 1× TBE buffer under 95V for 1:40 h.

Alleles were scored as presence or absence (1 or 0) using GelJ v2.0 (Heras et al., 2015). Population genetic structure was analyzed using STRUCTURE v2.3.4 (Pritchard et al., 2000) and the package pophelper (Francis, 2017). Genetic variation among populations was examined using a principal component analysis (PCA) in R.

### Statistical analyses

#### Microbiome richness and environmental and genetic correlates

To test whether northern duckweed populations harbor more bacterial richness than southern populations, we conducted a general linear mixed model (LMM) with region (northern vs. southern) as the predictor and a nested random effect (states nested within regions) using the package lme4 (Bates et al., 2015). We conducted the LMM for both observed ASV richness and asymptotic ASV richness (Chao estimator) using iNEXT. To identify which environmental factors might influence the geographic distribution of bacterial richness, we focused on 19 climatic and 13 water chemistry variables. We extracted the 19 climatic variables (WorldClim v2.1, 1970–2000) at 30 arc second resolution for the 34 populations. For water chemistry variables, we focused on pH, EC, TDS, nutrients (TOC, TN, TP, and C/N carbon to nitrogen ratio), and major and trace elements (Na, Ca, Mg, Si, Fe, and Mn). We did not consider some trace elements (Cd, Cu, Pb, and Zn) that showed little variation among populations or below the detection level (0.001 mg/L, Table S1). The water chemistry variables (except pH) were natural log transformed (log (x+0.01)) for analyses. For the climatic or water chemistry variables, we first conducted univariate regressions (general linear models, LMs) to select potential candidate predictors to be included in multiple regressions. We then used stepwise model selection (i.e. both forward and backward selections) of the multiple regressions based on the Akaike Information Criterion (AIC) to select the most parsimonious model and identify significant predictors. The lack of collinearity was confirmed based on the variance inflation factor (VIF). Duckweed genetic variation, represented by the first two axes of the genetic PCA (genetic PC1 and genetic PC2; Fig. 1c), was identified as non-significant predictors of bacterial richness by univariate regressions.

#### Microbiome composition and environmental and genetic correlates

To examine how diverse ecological processes, such as niche-based selection (by environments and host genetics), dispersal limitation, and chance, shaped bacterial community composition, we conducted four analyses. First, to assess the degree of distance decay in bacterial community similarity, we conducted a Mantel test between bacterial communities (the Bray–Curtis distance) and geographic distance using the package vegan (Oksanen et al., 2022). We further examined whether such distance decay was explained by geographic distance alone or environments. To do so, we conducted partial Mantel tests for climatic distance (all 19 climatic variables) and for water chemistry distance (all 13 water chemistry variables) while controlling for geographic distance. The climatic and water chemistry variables were (z-score) standardized prior to the estimation of their Euclidean distance among populations. The geographic distance was estimated based on the latitudes and longitudes of the populations (Table S1) using the package geodist (Padgham, 2021). Second, to quantify the relative importance of selection, dispersal, and chance in driving microbiome assembly among populations, we used a phylogenetic binning based null model analysis (iCAMP, Ning et al., 2020). Third, to further identify which environmental variables contributed to selection, we conducted univariate constrained principal component analysis (cPCoA) to select for potential predictors that may influence bacterial community composition. For the climatic variables, univariate cPCoAs revealed significant impact of all the 19 climatic variables, and thus we used the first two axes of the PCA of these climatic variables (climatic PC1 and PC2, accounting for 72.4% and 17.6% of total variation, respectively; Fig. S2). For water chemistry, univariate cPCoAs identified the impact of seven variables (TN, TP, C/N, Ca, Mg, Fe, and pH), and we further used multivariate cPCoAs and stepwise model selection to reduce the potential water chemistry predictors to be included together with climatic PC1 and climatic PC2 for final model selection. The lack of collinearity was confirmed using VIFs. Fourth, to examine the influence of duckweed genetic variation, which can be potentially shaped by environmental selection (see analysis below), on bacterial community composition, we conducted variation partitioning of bacterial communities using the package vegan among duckweed genetic variation (genetic PC1 and genetic PC2), climate and water chemistry (with predictors identified by model selections described above).

#### Duckweed genetic variation and environmental correlates

To examine how duckweed genetic variation was influenced by environments, we used univariate and multiple regressions with stepwise model selection to identify significant environmental predictors of genetic PC1 and genetic PC2. As univariate regressions revealed significant impacts of many climatic variables on genetic PC1 and genetic PC2, we used climatic PC1 and climatic PC2 as potential predictors, along with the water chemistry predictors identified by univariate regressions, in multiple regressions for model selection.

## Results

### Duckweed populations and microbiomes

Similar to terrestrial plants (Wei & Ashman, 2018; Acosta et al., 2020), duckweed microbiomes were dominated by Proteobacteria (79% of the ASVs), especially Alphaproteobacteria (42%) and Gammaproteobacteria (36%), followed by Bacteroidetes (7%), Actinobacteria (5%), Firmicutes (3%), and others (Fig. 1b). The microbiomes of duckweeds collected from the same populations (MA, Fig. 1b) were similar regardless of the sampling time (either during the peak or at the end of the growing season). Our analysis of duckweed genetic data revealed evidence of admixture (Fig. S3). We observed genetic differentiation between northern and southern populations along both the genetic PC1 and PC2 (Fig. 1c). We further found that genetic variation among duckweed populations was influenced by climate and water chemistry (Table S3). Specifically, duckweed genetic PC1 was influenced by precipitations (climatic PC2; multiple regression, LM: *t* = 3.57, *P* = 0.002) and water TN (*t* = 2.26, *P* = 0.035), and marginally by pH (*t* = -1.96, *P* = 0.063; Table S3). Duckweed genetic PC2 was primarily influenced by temperatures (climatic PC1, *t* = 5.80, *P* < 0.001; Table S3).

### Geographic variation of duckweed microbiome richness

To test whether bacterial richness is higher in northern duckweed populations compared to southern populations, we used a LMM and found that the northern populations hosted 10% more bacterial ASVs than the southern populations (LS-mean; observed richness: northern = 350 ± 30, southern = 321 ± 28, Fig. 2a; asymptotic richness: northern = 428 ± 44; southern = 388 ± 39; Fig. S4), while mean difference between northern and southern populations was not statistically significant (*P* > 0.05; Fig. 2a and Fig. S4).

**Fig. 2.**
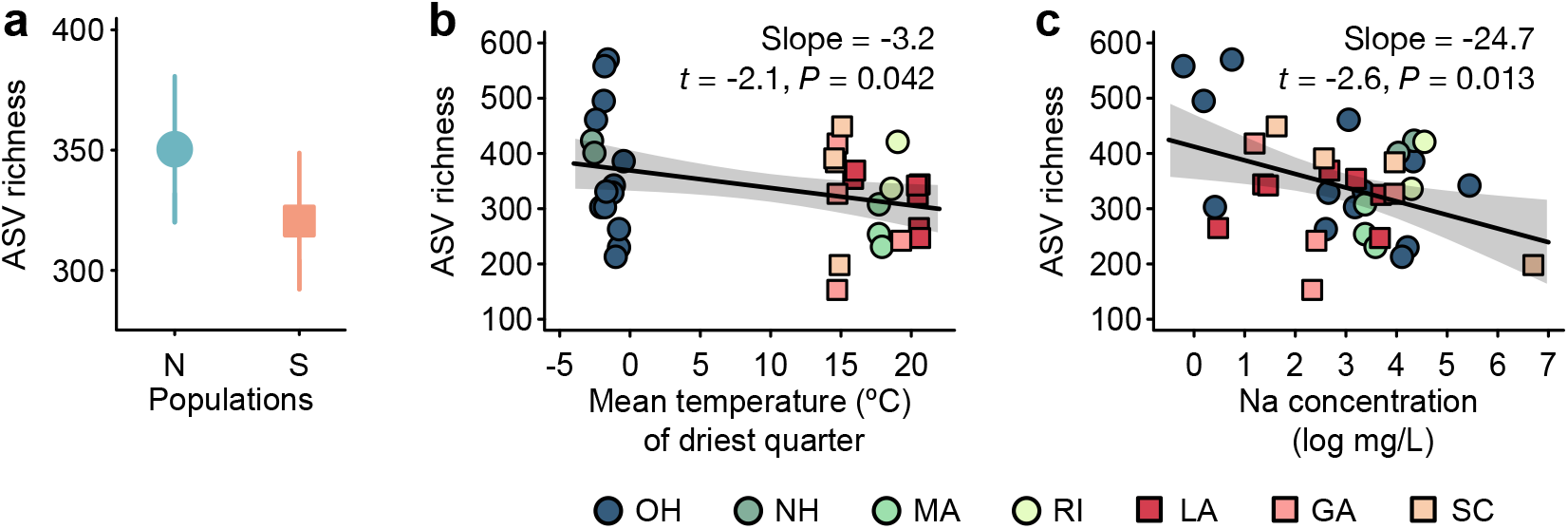
Environmental drivers of *Lemna* microbiome richness. (a) The least-squares mean (LS mean) ± SE of bacterial ASV richness are plotted for the northern populations (‘N’: OH, NH, MA, RI) and southern populations (‘S’: LA, GA, SC) using a general linear mixed model with region (northern vs. southern) as the predictor and states nested within regions as the random effect. (b) The mean temeprature of driest quarter (BIO9) and (c) the (natural log transformed) Na concentration of aquatic environments were identified as the important factors driving the distribution of bacterial richness of *Lemna* microbiomes after model selection of multiple regressions. Slopes with shaded 95% confidence intervals are shown. For statistical details, see Table S4.

Among the 19 climatic variables, only the mean temperature of the driest quarter (BIO9) showed a significant impact on bacterial richness, with a negative association observed between temperature and bacterial richness (multiple regression, LM: *t* = -2.12, *P* = 0.042; Fig. 2c and Fig. S4; Table S4). For water chemistry, while both concentrations of Na and TP were identified as potential factors influencing duckweed bacterial richness by univariate regressions, the multiple regression revealed that only Na concentration had a significant impact on bacterial richness, with lower richness associated with higher Na concentrations (LM: *t* = -2.63, *P* = 0.013; Fig. 2c and Fig. S4; Table S4). Unlike climate and water chemistry, the genetic variation of duckweed populations (genetic PC1 and PC2) did not influence bacterial richness (*P* > 0.05; Table S4).

### Geographic variation of duckweed microbiome composition

Duckweed bacterial communities exhibited distance decay in similarity (Mantel test, *r* = 0.46, *P* = 0.001; Fig. 3a). Such distance decay was not solely driven by geographic distance, but also by environmental factors (*r*_Climate|Geo_ = 0.27, *P* = 0.001; *r*_Water chemistry|Geo_ = 0.29, *P* = 0.001). This result indicated that both selection and dispersal limitation as well as chance influenced duckweed microbiome assembly. We further found that selection played an important role (26%) in structuring duckweed bacterial communities, in addition to dispersal limitation (33%) and chance (and other unidentified weak processes, 41%; Fig. 3b).

**Fig. 3.**
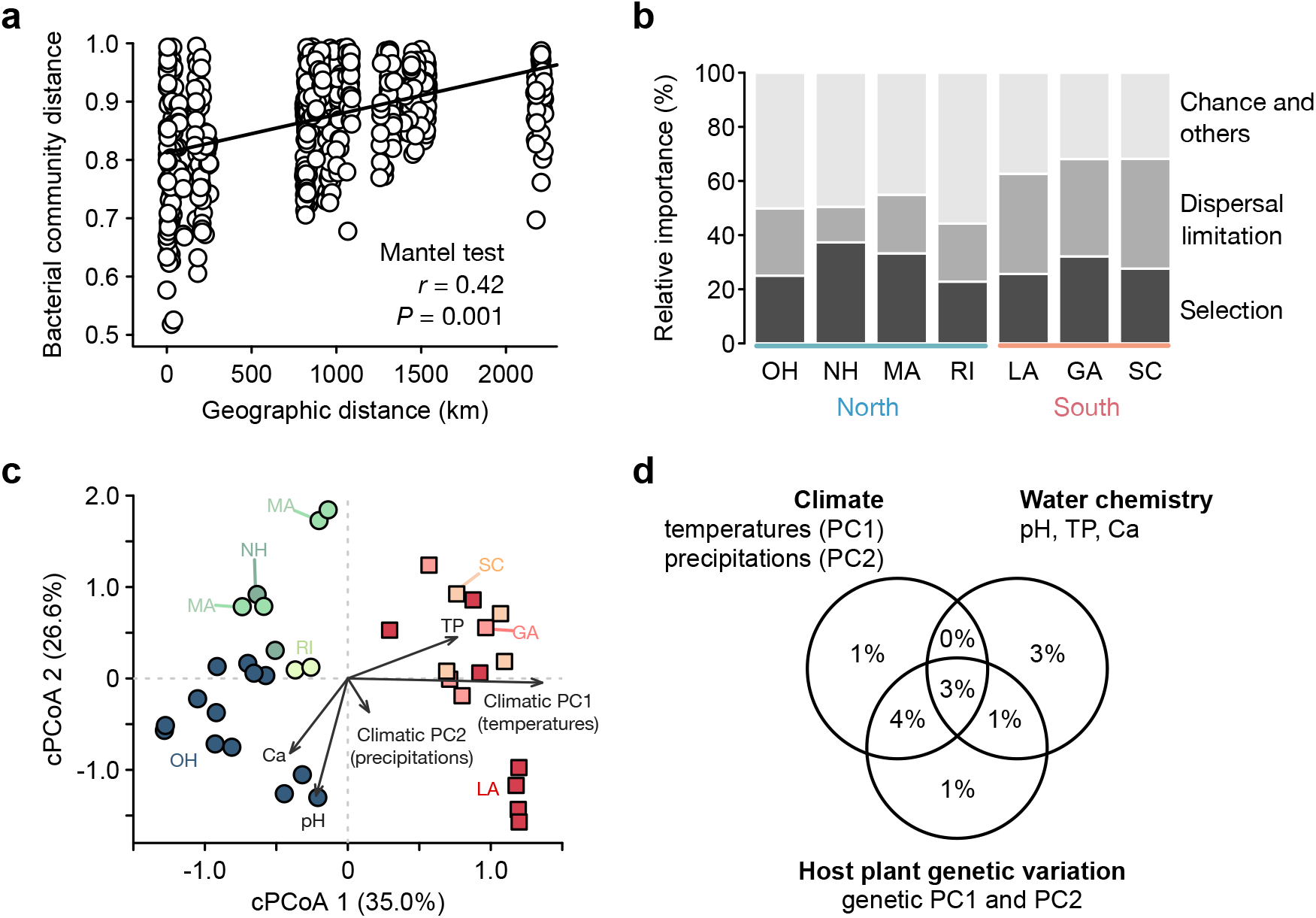
Ecological processes underlying the geographic variation of *Lemna* microbiome composition. (a) The Mantel test indicates a significant correlation between the Bray-Curtis distance of bacterial communities and geographic distance. (b) The relative importance of ecological processes driving *Lemna* microbiome assembly was quantified using the package iCAMP. We focused on selection (homogeneous and heterogeneous selections), dispersal limitation, and chance, whereas ‘others’ encompass weak processes (iCAMP) including homogenizing dispersal here. (c) The first two axes of the 19 climatic variables (climatic PC1 and climatic PC2), pH, and concentrations of total phosphorus (TP) and calcium (Ca) were identified as the most important factors driving variation in *Lemna* bacterial community composition after model selection of constrained principal component analyses (cPCoAs). (d) Variation partitioning indicates the collective roles of duckweed genetic variation, climate, and water chemistry in explaining the geographic variation of *Lemna* bacterial community composition. For statistical details, see Table S5.

Among the environmental factors, climatic PC1 (temperatures) and PC2 (precipitations) together with water pH, TP, and Ca were the most important variables driving the geographic variation of duckweed bacterial community composition (cPCoA: climatic PC1, 7.2% of variation, *F* = 2.9, *P* = 0.001; climatic PC2, 4.3%, *F* = 1.7, *P* = 0.006; pH, 5.7%, *F* = 2.3, *P* = 0.001; TP, 3.6%, *F* = 1.4, *P* = 0.048; Ca, 3.9%, *F* = 1.6, *P* = 0.012; Fig. 3c and Table S5). Climatic PC1 (temperatures) and TP were found to influence bacterial community cPCoA 1, while climatic PC2 (precipitations), pH, and Ca were found to influence cPCoA 2 (Fig. 3c). Based on the subset of populations (*N* = 25) with duckweed genetic data, we found that duckweed genetic variation affected bacterial community composition (cPCoA: genetic PC1, 7.7%, *F* = 2.0, *P* = 0.001; genetic PC2, 9.9%, *F* = 2.6, *P* = 0.001; Table S5). Variation partitioning analysis further pointed out the collective roles of climate, water chemistry, and host genetic variation on duckweed bacterial community composition (Fig. 3d).

## Discussion

Our study on the microbiomes of wide-ranging duckweeds revealed that the geographic distribution of plant microbiome diversity supported the standing hypothesis of microbial biogeography, with bacterial richness higher in temperate regions relative to lower latitudes as observed in free-living environmental microbiomes. We also found that temperature (of the driest quarter, BIO9) and Na concentration showed a negative impact on the distribution of duckweed bacterial richness, while host genetic variation showed no strong effect. In contrast to bacterial richness, the geographic variation of duckweed bacterial community composition was influenced by all 19 climatic variables, including temperatures (climatic PC1) and precipitations (climatic PC2), and water chemistry variables such as pH and concentrations of TP and Ca. Our results further underscored the collective roles of host genetic variation, climate, and water chemistry in driving duckweed bacterial community composition.

### Bacterial richness of plant microbiomes is higher in temperate populations

Our findings of higher bacterial richness in temperate relative to subtropical duckweed populations were consistent with global patterns of microbial biogeography in free-living microbiomes across ecosystems, including soil, marine, and airborne microbiomes (Tedersoo et al., 2014; Sunagawa et al., 2015; Bahram et al., 2018; Zhao et al., 2022). Similar to wild plants such as duckweeds, a latitudinal pattern of increased bacterial richness has also been observed in crops such as the rhizosphere microbiomes associated with soybean from tropical to temperate regions (Zhang et al., 2018). In our study, we observed a 10% higher bacterial richness in temperate duckweed populations compared to subtropical populations, while the mean difference between the two regions was not statistically significant. This suggests that other factors, which do not follow a latitudinal pattern, might influence duckweed bacterial richness, such as Na concentration in freshwater ecosystems (Fig. 2c). We found that Na concentration negatively impacted bacterial richness in these natural duckweed populations. This negative impact of Na concentration on microbial growth has also been demonstrated experimentally in duckweeds (O’Brien et al., 2020). Interestingly, we observed high Na concentration in some populations from both temperate and subtropical regions (Table S1), potentially reflecting road salt use in the north and proximity to seawater in the south. This suggests that factors such as increased salinity in freshwater ecosystems due to, for instance, road salt flux (Kaushal et al., 2005; Hintz et al., 2022) and sea level rise (Jackson & Jevrejeva, 2016; Dangendorf et al., 2017), as well as increased temperature (IPCC, 2022), under global change may have negative impacts on plant microbiome richness and their latitudinal patterns.

### Environmental factors influence the geographic variation of plant microbiome composition

The geographic variation of duckweed bacterial community composition among populations exhibited distance decay, driven by diverse ecological processes. Among these processes, dispersal limitation and chance played a major role (74%), similar to the observations (70–80%) in global distributions of free-living soil and marine microbiomes (Zhao et al., 2022). Consistent with global soil microbiomes (Zhao et al., 2022), selection accounted for 26% of the influence in driving the geographic variation of duckweed bacterial community composition. Specifically, environmental pH, which is a dominant driver of global soil bacterial community composition (Fierer & Jackson, 2006; Bahram et al., 2018), was also found to influence duckweed bacterial community composition in the aquatic environments here. Similar to marine microbiomes (Sunagawa et al., 2015), temperatures strongly impacted duckweed bacterial community composition. Such effects of temperature and pH on bacterial community composition have also been demonstrated experimentally in duckweeds (Calicioglu et al., 2018). Additionally, phosphorus, one of the most important limiting factors in freshwater ecosystems (Hudson et al., 2000), influenced duckweed bacterial community composition, similar to observations in bacterial communities associated with marine algae (Martin et al., 2021). Furthermore, we found that calcium concentration, reflecting hardness of aquatic environments, was also driving duckweed bacterial community composition, independent from the strong impact of pH (after model selection). Our study, together with previous research, point to some general principles of microbial biogeography regarding the influence of selection by environments and the underlying drivers. These findings highlight the potential impacts on the distribution of microbiome composition of climate change and anthropogenic activities, particularly in terms of nutrient deposition and discharge into ecosystems (Schlesinger, 2009; Tipping et al., 2014), and the overall quality of aquatic environments.

### Host genetic variation plays a role in the geographic distribution of plant microbiomes

While we have highlighted the similarities in the patterns and mechanisms of microbial biogeography between plant microbiomes and free-living environmental microbiomes, our findings also emphasized the joint role of plant genetic variation and environmental variation. Different from soil microbiomes, where aboveground plant diversity does not influence bacterial community composition (Fierer & Jackson, 2006; Tedersoo et al., 2014; Bahram et al., 2018), our study showed that plant genetic variation influenced duckweed bacterial community composition (Table S5) via its joint effect with climate and water chemistry, rather than their independent effects (Fig. 3d). This was primarily because the genetic variation of duckweeds was strongly influenced by the same factors that influenced their microbiome composition, such as temperatures, precipitations, nitrogen concentration (which was correlated with phosphorus concentration), and pH. The strong coupling of host genetic variation and microbiomes with environmental factors made it challenging to separate the effects of host genetic and environmental variation on microbiome composition in natural populations without manipulative experiments. This observation should not be unique to duckweeds but is expected to be common in plant microbiomes, because local adaptation to environments is a widespread phenomenon in plants (Leimu & Fischer, 2008). This observation underscores the potential for even stronger impacts on the distribution, structure, and function of plant microbiomes in the cases of misaligned responses between plants and microbes to climate change and anthropogenic activities.

## Conclusions

Our study elucidates the geographic distribution of plant microbiome structure and the underlying mechanisms, highlighting both the commonalities and differences in microbial biogeography relative to free-living environmental microbiomes. Our findings call for the need of additional research across diverse plant species and populations, geographic scales, and ecosystems to further advance our understanding of the principles of microbial biogeography. The key drivers identified in our study, including temperatures, precipitations, pH, and concentrations of sodium, phosphorus, and calcium, along with host genetic variation, provide important insights into predicting the vulnerability and resilience of plant microbiomes and their impacts on ecosystem functioning under changing climates and intensifying anthropogenic activities.

## Acknowledgements

We thank Maris Hollowell for the assistance in microbiome DNA extraction and duckweed genotyping, Jada Daniels, Zhijun Ding, Jessica LaBella, Cordelia Zheng, Olivia Walker, and Reetu Shrestha for the assistance in sample collection and processing.

---

**Fig. S1.**
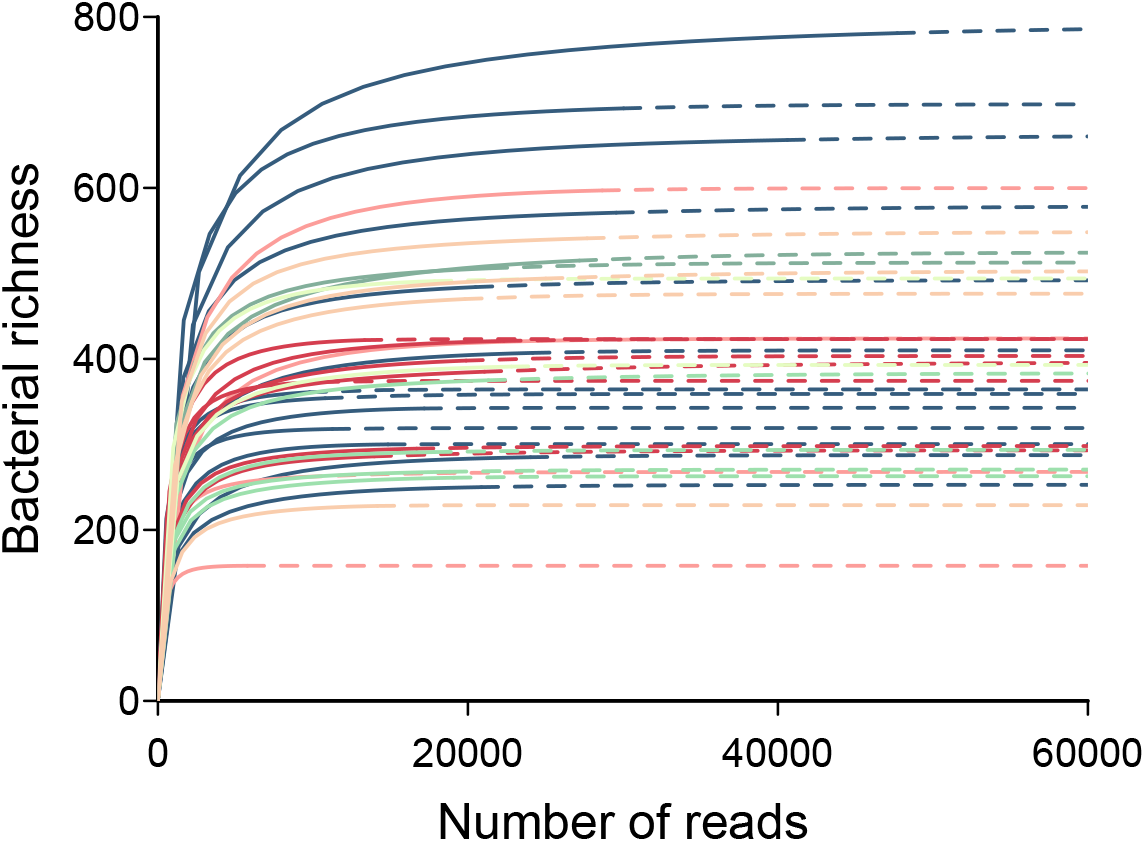
Rarefaction reveals that the majority of the bacterial richness of *Lemna* microbiomes was captured by the sequencing effort. The number of reads is represented by the solid portion of each line, whereas the dashed portion indicates extrapolation in the rarefaction analysis using the R package iNEXT. Colors indicate the different origins (states) of duckweed populations.

**Fig. S2.**
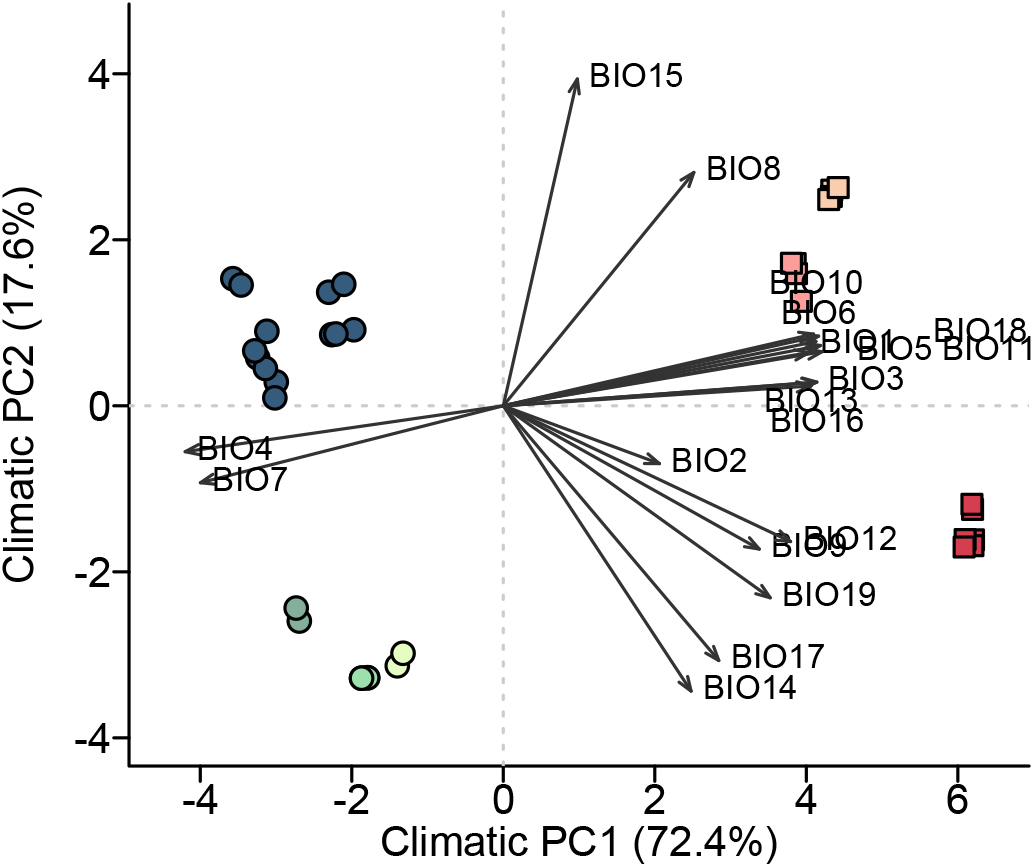
The climatic PCA of *Lemna* populations. The climatic PCA was based on the 19 climatic variables of the 34 *Lemna* populations. BIO1 = Annual Mean Temperature; BIO2 = Mean Diurnal Range (Mean of monthly (max temp - min temp)); BIO3 = Isothermality (BIO2/BIO7) (×100); BIO4 = Temperature Seasonality (standard deviation ×100); BIO5 = Max Temperature of Warmest Month; BIO6 = Min Temperature of Coldest Month; BIO7 = Temperature Annual Range (BIO5-BIO6); BIO8 = Mean Temperature of Wettest Quarter; BIO9 = Mean Temperature of Driest Quarter; BIO10 = Mean Temperature of Warmest Quarter; BIO11 = Mean Temperature of Coldest Quarter; BIO12 = Annual Precipitation; BIO13 = Precipitation of Wettest Month; BIO14 = Precipitation of Driest Month; BIO15 = Precipitation Seasonality (Coefficient of Variation); BIO16 = Precipitation of Wettest Quarter; BIO17 = Precipitation of Driest Quarter; BIO18 = Precipitation of Warmest Quarter; BIO19 = Precipitation of Coldest Quarter.

**Fig. S3.**
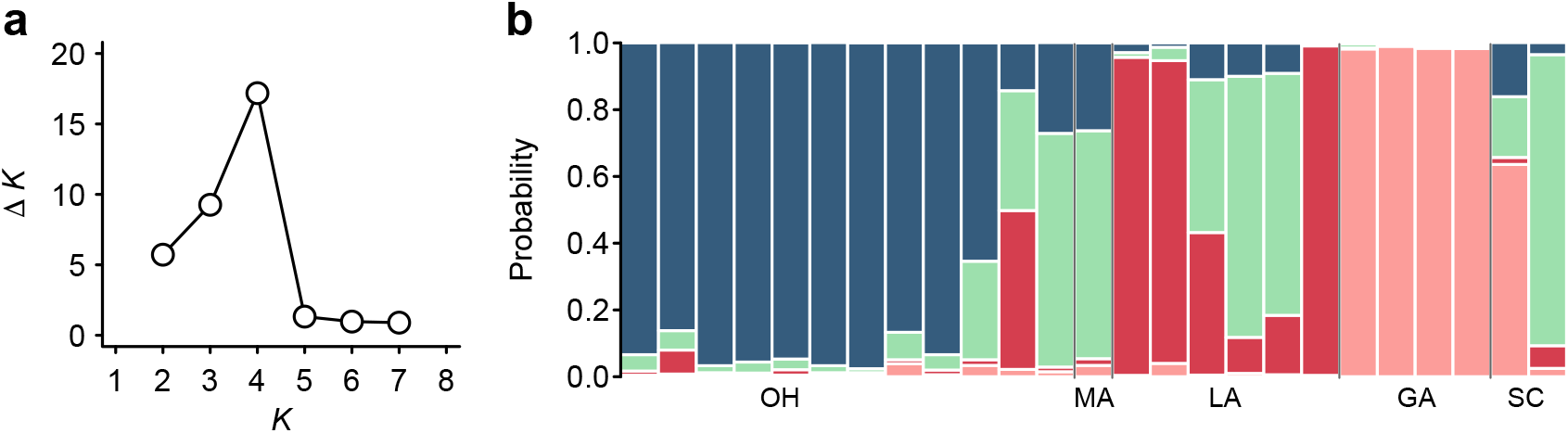
STRUCTURE results of *Lemna*. (a) The inference of populations (*K*) identifies four genetic clusters. (b) Inferred admixture plot of the 25 *Lemna* samples is displayed at *K* = 4.

**Fig. S4.**
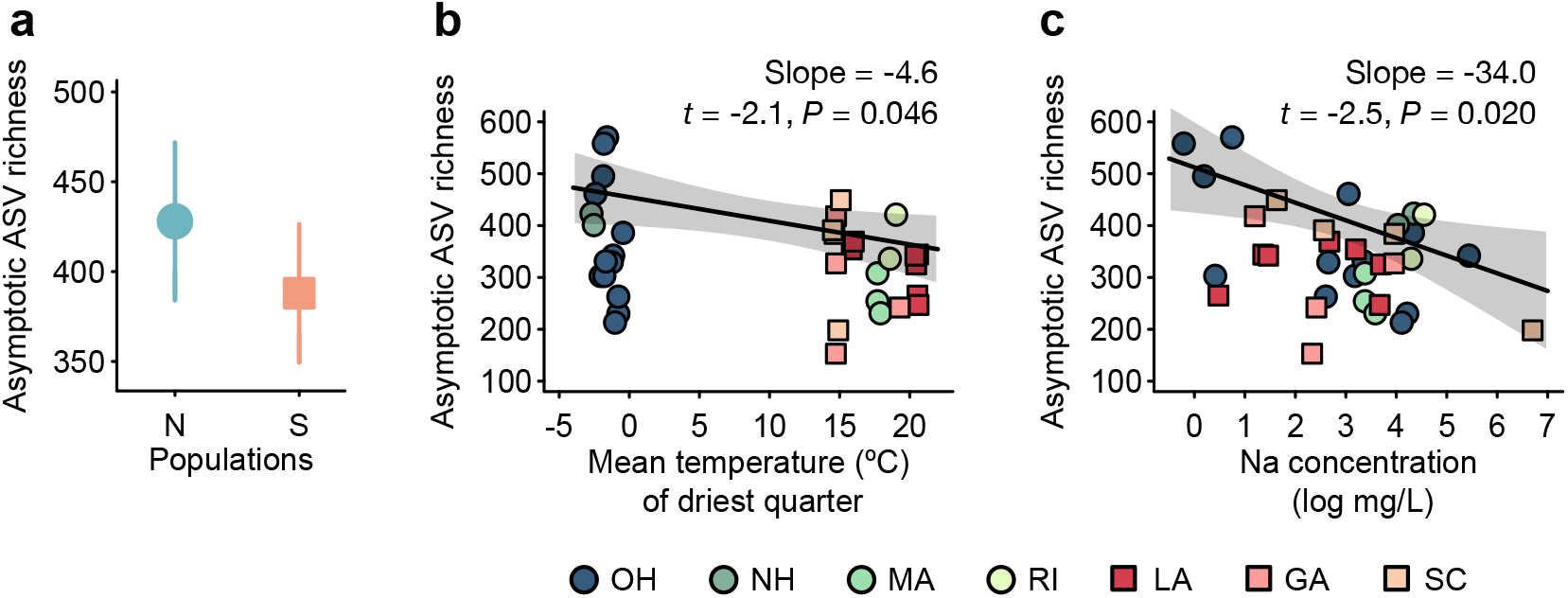
Environmental drivers of the asymptotic richness of *Lemna* bacterial microbiomes. (a) The least-squares mean (LS mean) ± SE of bacterial ASV richness (based on Chao estimator) are plotted for the northern populations (OH, NH, MA, RI) and southern populations (LA, GA, SC), using a general linear mixed model with region (northern vs. southern) as the predictor and states nested within regions as the random effect. (b) The mean temeprature of the driest quarter (BIO9) and (c) the (natural log transformed) Na concentration of aquatic environments were identified as the important factors driving the distribution of *Lemna* bacterial richness after model selection of multiple regressions. Slopes and shaded 95% confidence intervals are shown. For statistical details, see Table S4.

